# An Information Theoretic Framework for Protein Activity Measurement

**DOI:** 10.1101/2021.10.02.462873

**Authors:** Aaron T. Griffin, Lukas J. Vlahos, Codruta Chiuzan, Andrea Califano

## Abstract

Nonparametric analytical Rank-based Enrichment Analysis (NaRnEA) is a novel gene set analysis method which leverages an analytical null model derived under the Principle of Maximum Entropy. NaRnEA critically improves over two widely used methods – Gene Set Enrichment Analysis (GSEA) and analytical Rank-based Enrichment Analysis (aREA) – as shown by differential activity measurement of ~2,500 transcriptional regulatory proteins across three cohorts in The Cancer Genome Atlas (TCGA) based on the enrichment of their transcriptional targets in differentially expressed genes. Phenotype-matched proteomic data from the Clinical Proteomic Tumor Analysis Consortium (CPTAC) was used to evaluate measurement accuracy. We show that the sample-shuffling empirical null models leveraged by GSEA and aREA are overly conservative, a shortcoming that is critically addressed by NaRnEA’s optimal analytical null model.

## Introduction

The ever-increasing availability of next-generation sequencing technologies and highly accurate annotations for prokaryotic and eukaryotic genomes, especially the human genome, have transformed biology into a data-rich scientific discipline^1^. Thus, leveraging large-scale, gene-level biochemical measurements to support mechanistic inferences involving biological and cellular processes^2,3^, as well as to measure protein and pathway activity^4^, has become a critical priority in computational biology^5^.

Not surprisingly, gene set analysis methods, which were developed precisely to address some of these challenges, have rapidly emerged as some of the most utilized tools in biomedical research. These methods provide a statistical assessment of the biological relevance of a specific set of genes – ranging from signal transduction pathways, gene ontology categories, and transcriptional targets of regulatory proteins, among others – based on their ranking according to a quantitative metric of interest. Most frequently, differential expression between two cellular states is used as the ranking criterion, but virtually any criteria to rank genes in a list can be used; see (Maleki et al. 2020 Front Genet)^6^ and (Das et al. 2020 Entropy)^7^ for recent reviews comparing the wide variety of published gene set analysis methods, as well as the statistical assumptions implicit to each method.

In this manuscript, we leverage the Principle of Maximum Entropy from Information Theory^8^ to derive a provably optimal solution for determining the statistical significance of gene set enrichment which is implemented in the novel algorithm Nonparametric analytical Rank-based Enrichment Analysis (NaRnEA). We show that NaRnEA overcomes key limitations of two widely used gene set analysis methods – Gene Set Enrichment Analysis (GSEA)^9^ and analytical Rank-based Enrichment Analysis (aREA)^4^. In addition, while the magnitude of a gene set’s enrichment test statistic is strongly dependent on the size of the gene set, NaRnEA provides a Proportional Enrichment Score (PES) as an effect size that is virtually independent of gene set size. For simplicity, we focus on enrichment analysis in the context of differential gene expression; however, NaRnEA can trivially generalize to other quantitative ranking metrics and datasets.

Accurate benchmarking of gene set analysis methods using experimental data is challenging because the gene sets commonly used by these methodologies are often literature-based lists (e.g. KRAS pathway genes^10^) that lack any kind of independent, systematic validation^5–7^. However, recent work in cancer systems biology has shown that these methods can also be used to accurately measure the differential activity of transcriptional regulatory proteins, which provides a more objective benchmark that can be experimentally assessed. We define differential protein activity as the contribution of a transcriptional regulatory protein to the implementation of a specific differential gene expression signature. Consistent with this definition and akin to using a highly multiplexed gene reporter assay, we introduced the Virtual Inference of Protein Activity by Enriched Regulon Analysis (VIPER) algorithm^4^ to measure protein activity based on the enrichment of transcriptional targets (i.e. regulon gene set) in a signature of differentially expressed genes.

The tissue-specific regulons necessary for these analyses can be effectively reverse-engineered using a variety of methods^11^. In the context of this study, we use the ARACNe3 algorithm, which is the most recent implementation of the Algorithm for the Reconstruction of Accurate Cellular Networks (ARACNe)^12–14^; the previous versions of this algorithm have been experimentally validated as ARACNE-inferred regulons have been used extensively to measure differential protein activity in combination with VIPER, effectively elucidating Master Regulator (MR) proteins representing mechanistic determinants of tumor transcriptional state^15–17^. This paradigm has also been extended to measure differential protein activity in single cells, even when no transcripts can be detected for the encoding gene, with accuracy comparable to antibody-based experimental methods^18^. Thus, large-scale, mass spectrometry-based protein abundance protein abundance measurements and phenotype-matched gene expression profiles provide a unique, objective benchmark to evaluate improvements to the measurement of differential protein activity using gene set analysis methodologies.

Our study shows that, when assessed across three primary tumor cohorts from TCGA – lung adenocarcinoma (LUAD), colon adenocarcinoma (COAD), and head-neck squamous cell carcinoma (HNSC) – NaRnEA measures statistically significant differential activity for hundreds of transcriptional regulatory proteins that is otherwise undetectable by GSEA and aREA. Crucially, CPTAC proteomic data analysis showed significant correlation between differentially active proteins, as measured by NaRnEA, and differential protein abundance, as measured from CPTAC data.

NaRnEA and ARACNe3 are freely available for research use on GitHub (https://github.com/califano-lab/NaRnEA).

## Results

### Nonparametric analytical Rank-based Enrichment Analysis

NaRnEA is a nonparametric, frequentist statistical method designed to infer gene set enrichment according to the paradigm of null hypothesis significance testing. The nonparametric formulation of NaRnEA enables inference of gene set enrichment in gene expression signatures estimated with a variety of differential gene expression methods (e.g. DESeq2^19^, Mann-Whitney U test). NaRnEA requires each member of a gene set to be parameterized with two values: (i) the Association Weight and (ii) the Association Mode. The Association Weight represents the likelihood that the specific gene truly belongs to the gene set. For example, in gene sets representing the regulon of a transcriptional regulatory protein, the Association Weight would indicate the confidence in the gene as a *bona fide* transcriptional target of the protein. The Association Mode represents whether the expression of the gene is positively or negatively associated with the enrichment of the gene set. This would, for instance, differentiate positive and negative regulators of apoptosis in a literature-curated apoptosis gene set. The Association Weight is a positive value, while the Association Mode should be a value in the [−1, 1] range. As a result, NaRnEA can assess enrichment of simple gene sets, such as those constructed based on biological processes or pathways, as well as enrichment of more complex gene sets that incorporate substantial prior knowledge, such as regulons whose genes are associated with a specific likelihood and regulation directionality.

NaRnEA derives a fully analytical null model for gene set enrichment using the information theoretic Principle of Maximum Entropy (see Methods). Leveraging this provably optimal null model, NaRnEA calculates a gene set enrichment test statistic – referred to as the Normalized Enrichment Score (NES) – which is an asymptotically normal random variable when the gene set is not enriched in the gene expression signature (i.e. the null hypothesis for gene set analysis). A statistically significant positive or negative value of the NaRnEA NES provides evidence for positive or negative gene set enrichment, respectively. In addition to the NES, NaRnEA calculates a gene set enrichment effect size – referred as the Proportional Enrichment Score (PES) (see Methods). The NaRnEA PES is calculated by adjusting the NaRnEA NES by its maximum (or minimum) possible value in the case of positive (or negative) gene set enrichment, thus accounting for features specific to the gene set’s size and parameterization as well as the gene expression signature itself. The NaRnEA PES is positive when the gene set is positively enriched, negative when the gene set is negatively enriched, and has a maximum magnitude of 1, thus providing a metric that is universally comparable between different gene set sizes (i.e. gene set size-independent) and across different gene expression signatures.

### Gene Set Analysis Method Comparison

GSEA requires properly normalized gene expression profiles from samples representing a test phenotype and a reference phenotype; the differential expression of each gene is then estimated using an appropriate statistical test (e.g. Mann-Whitney U test). The enrichment of a gene set in this gene expression signature is calculated with a weighted Kolmogorov-Smirnov-like statistic (i.e. the GSEA enrichment score). It is recommended that the null distribution of the GSEA enrichment score for a particular gene set should be approximated using an empirical phenotype-based permutation procedure^9,20^; two-sided empirical p-values are calculated accordingly.

aREA accepts the same input data with differential gene expression estimated using Welch’s unpaired t-test. The enrichment of a gene set in this gene expression signature is calculated using a three-tailed approach, returning an enrichment score test statistic. It is recommended that the null distribution for the aREA enrichment score for a particular gene set should also be approximated using an empirical phenotype-based permutation procedure or an approximate analytical approach if there are not enough samples for the permutation-based strategy; two-sided empirical p-values are calculated accordingly^4^.

### Benchmarking Gene Set Analysis Methods based on Differential Protein Activity Measurement

To compare the performance of different gene set analysis methods, we used NaRnEA, GSEA, and aREA to measure the differential activity of 2,491 transcription factor (TF) and co-transcription factor (coTF) proteins based on the enrichment of their ARACNe3-inferred regulons (i.e. transcriptional target gene sets) in genes differentially expressed between primary tumor samples and adjacent normal tissue samples in three cohorts from TCGA as discussed in (*Alvarez et al. Nat Genet 2016*)^4^. For this, we reverse-engineered a tumor-specific regulons (TS-regulons) for each TF and coTF using LUAD, COAD, or HNSC primary tumor gene expression profiles from the respective TCGA cohorts. These cancer types were selected because of (a) the availability of phenotype-matched primary tumor and adjacent normal tissue RNA-Seq gene expression profiles in TCGA and (b) the availability of phenotype-matched primary tumor and adjacent normal tissue mass spectrometry protein abundance profiles in CPTAC, supporting direct assessment of transcriptional regulatory proteins that have differential activity and differential abundance between malignant and normal tissues. To avoid potential overfitting, primary tumor gene expression profiles which were used in the calculate of differential gene expression signatures were excluded from regulon reverse-engineering with ARACNe3. The remaining gene expression profiles from each cohort were used to reverse-engineer a TS-regulon for each TF and coTF protein with ARACNe3, the latest implementation of the ARACNe algorithm (see Methods). A null model regulon (NM-regulon) was constructed from each TS-regulon by replacing ARACNe3-inferred transcriptional targets with an equal number of genes randomly selected from genes not originally in the TS-regulon (i.e. the regulon gene set complement).

The LUAD transcriptional regulatory network was reverse-engineered from 476 primary tumor gene expression profiles and contains 2,491 regulators, 19,351 targets, and 947,776 interactions. The COAD transcriptional regulatory network was reverse-engineered from 437 primary tumor gene expression profiles and contains 2,491 regulators, 19,351 targets, and 646,341 interactions. The HNSC transcriptional regulatory network was reverse-engineered from 457 primary tumor gene expression profiles and contains 2,491 regulators, 19,351 targets, and 810,683 interactions.

Based on previous work, we assume that enrichment of TS-regulons in a differential gene expression signature will effectively measure the differential activity of the corresponding transcriptional regulatory protein^4,18^, while NM-regulons should not produce any statistically significant enrichment, thus providing an optimal comparative null model. It is thus reasonable to expect that a subset of TS-regulons, yet no NM-regulons, will demonstrate statistically significant enrichment in the corresponding gene expression signature, as assessed by comparing gene expression profiles from primary tumor and adjacent normal tissue. In this benchmarking, TS-regulons which are not correctly inferred will behave like NM-regulons and will therefore not contribute to the difference in performance between the gene set analysis methods. We thus propose that the performance of a gene set analysis method can be evaluated based on (1) the ability of each method to differentiate between TS-regulons and NM-regulons, (2) the number of TS-regulons with statistically significant differential protein activity, (3) the number of NM-regulons with statistically significant differential protein activity, and (4) the agreement between the differential protein activity measured by each method and the differential protein abundance measured by mass spectrometry.

### Gene Set Analysis Sensitivity and Specificity

We used NaRnEA, GSEA, and aREA to compute the enrichment of both TS-regulons and NM-regulons in differential gene expression signatures as assessed by comparing gene expression profiles of 57 LUAD, 41 COAD, and 43 HNSC patient-matched primary tumor and adjacent normal tissue samples from TCGA.

We generated receiver operating characteristic (ROC) curves for NaRnEA, GSEA, and aREA within each TCGA cohort by designating TS-regulons as cases (i.e. positives) and NM-regulons as controls (i.e. negatives) under the assumption that none of the NM-regulons should produce statistically significant enrichment, while a substantial fraction of the TS-regulons should be statistically significantly enriched in the corresponding tumor vs. tissue gene expression signature. The statistical significance of the Area Under the ROC curve (AUROC) was calculated using a one-tailed Mann-Whitney U test and the 95% confidence interval for each AUROC was estimated using DeLong’s method^21^; DeLong’s test was used to compare AUROC values for each pair of methods within each cancer subtype.

Based on this analysis, NaRnEA statistically significantly outperformed GSEA and aREA in its ability to differentiate between TS-regulons and NM-regulons based on the enrichment two-sided p-values, achieving a highly statistically significant AUROC in LUAD (AUROC = 0.741, p = 4.13e-191). COAD (AUROC = 0.698, p = 2.05e-129), and HNSC (AUROC = 0.716, p = 4.42e-154) (Figure 1). The AUROC for GSEA was statistically significant only in HNSC (AUROC = 0.523, p = 2.75e-3), while the AUROC for aREA was statistically significant only in LUAD (AUROC = 0.555, p = 1.07e-11) and HNSC (AUROC = 0.521, p = 5.71e-3). DeLong’s test shows that NaRnEA statistically significantly outperformed both GSEA and aREA in all three cohorts (FWER < 0.05).

**Figure 1.**
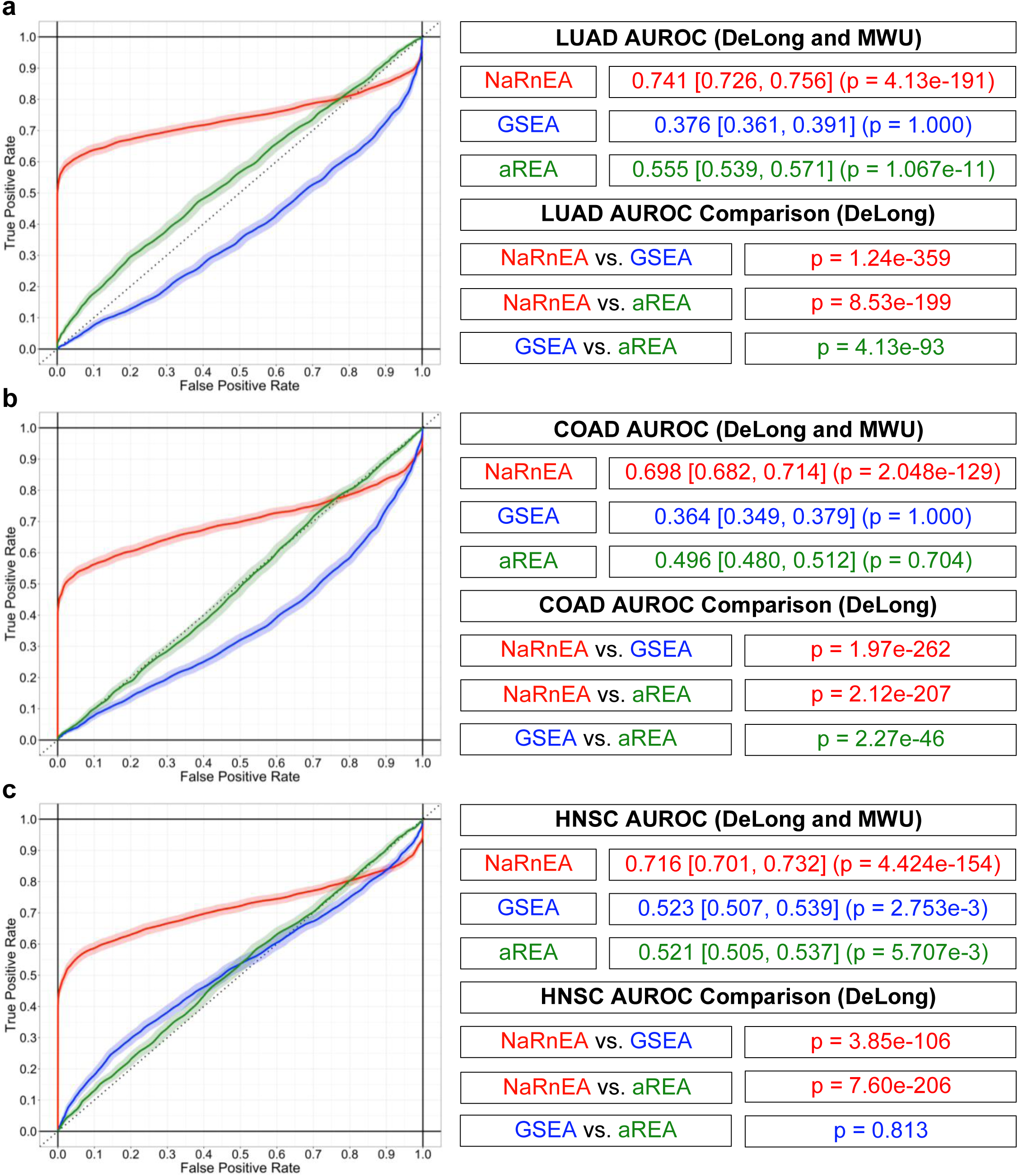
Comparative Benchmark of NaRnEA, GSEA, and aREA for Measurement of Differential Protein Activity. NaRnEA significantly outperforms GSEA and aREA for the purpose of distinguishing between TS-regulons and NM-regulons in TCGA transcriptomic data from (a) LUAD, (b) COAD, and (c) HNSC cancer cohorts. ROC curves are constructed from gene set enrichment two-sided p-values by setting ARACNe3-inferred regulon gene sets as cases and null gene sets as controls. 95% confidence intervals for ROC curves are estimated using 2,000 stratified bootstraps. 95% confidence intervals for the AUROC are calculated using DeLong’s method. The statistical significance of the AUROC is calculated with a one-tailed Mann-Whitney U (MWU) test. Within each cancer type, AUROC values for each method are compared in a pairwise manner using DeLong’s test. NaRnEA ROC curves are shown in red, GSEA ROC curves are shown in blue, and aREA ROC curves are shown in green.

Direct analysis of the two-sided p-values provides an estimate of the fraction of TS-regulons and NM-regulons identified as statistically significantly enriched by the different gene set analysis methods based on control of the False Positive Rate (FPR < 0.05), the False Discovery Rate (FDR < 0.05), or the Family Wise Error Rate (FWER < 0.05). In agreement with the ROC curve analysis, NaRnEA was the only gene set analysis method capable of identifying statistically significantly enriched TS-regulons after correcting for multiple hypothesis testing within each cancer cohort (Table 1). Furthermore, after correcting for multiple hypothesis testing, NaRnEA did not identify any statistically significantly enriched NM-regulons, demonstrating that NaRnEA is substantially more sensitive than GSEA or aREA, without loss of specificity.

**Table 1.** Comparative Analysis of NaRnEA, GSEA, and aREA. Comparing NaRnEA, GSEA, and aREA gene set enrichment two-sided p-values without correction (FPR < 0.05), after correcting to control the False Discovery Rate (FDR < 0.05), and after correcting to control the Family Wise Error Rate (FWER < 0.05). P-values were corrected for multiple hypothesis testing to control the FDR according to the methodology of Benjamini and Hochberg. P-values were corrected for multiple hypothesis testing to control the FWER according to the methodology of Bonferroni. Maximum likelihood estimates of the proportion of significantly enriched gene sets are shown as percentages along with 95% confidence intervals for the proportion which are calculated according to the procedure of Clopper and Pearson.

| Subtype | Gene Sets | Threshold | NaRnEA<br>[95% CI] | GSEA<br>[95% CI] | aREA<br>[95% CI] |
| --- | --- | --- | --- | --- | --- |
| LUAD | ARACNe3 | FPR < 0.05 | 60.98%<br>[59.03%, 62.90%] | 7.47%<br>[6.47%, 8.57%] | 9.39%<br>[8.28%, 10.61%] |
|  |  | FDR < 0.05 | 59.37%<br>[57.41%, 61.31%] | 0.00%<br>[0.00%, 0.15%] | 0.00%<br>[0.00%, 0.15%] |
|  |  | FWER < 0.05 | 44.92%<br>[42.96%, 46.90%] | 0.00%<br>[0.00%, 0.15%] | 0.00%<br>[0.00%, 0.15%] |
|  | Null | FPR < 0.05 | 4.94%<br>[4.12%, 5.86%] | 9.80%<br>[8.66%, 11.03%] | 4.05%<br>[3.31%, 4.91%] |
|  |  | FDR < 0.05 | 0.00%<br>[0.00%, 0.15%] | 0.00%<br>[0.00%, 0.15%] | 0.00%<br>[0.00%, 0.15%] |
|  |  | FWER < 0.05 | 0.00%<br>[0.00%, 0.15%] | 0.00%<br>[0.00%, 0.15%] | 0.00%<br>[0.00%, 0.15%] |
| COAD | ARACNe3 | FPR < 0.05 | 53.31%<br>[51.33%, 55.29%] | 6.14%<br>[5.23%, 7.16%] | 4.34%<br>[3.57%, 5.21%] |
|  |  | FDR < 0.05 | 50.82%<br>[48.84%, 52.80%] | 0.00%<br>[0.00%, 0.15%] | 0.00%<br>[0.00%, 0.15%] |
|  |  | FWER < 0.05 | 32.52%<br>[30.68%, 34.40%] | 0.00%<br>[0.00%, 0.15%] | 0.00%<br>[0.00%, 0.15%] |
|  | Null | FPR < 0.05 | 5.06%<br>[4.23%, 5.99%] | 7.91%<br>[6.88%, 9.04%] | 4.09%<br>[3.35%, 4.95%] |
|  |  | FDR < 0.05 | 0.00%<br>[0.00%, 0.15%] | 0.00%<br>[0.00%, 0.15%] | 0.00%<br>[0.00%, 0.15%] |
|  |  | FWER < 0.05 | 0.00%<br>[0.00%, 0.15%] | 0.00%<br>[0.00%, 0.15%] | 0.00%<br>[0.00%, 0.15%] |
| HNSC | ARACNe3 | FPR < 0.05 | 55.28%<br>[53.30%, 57.24%] | 5.78%<br>[4.90%, 6.77%] | 6.54%<br>[5.60%, 7.59%] |
|  |  | FDR < 0.05 | 52.47%<br>[50.49%, 54.45%] | 0.00%<br>[0.00%, 0.15%] | 0.00%<br>[0.00%, 0.15%] |
|  |  | FWER < 0.05 | 33.80%<br>[31.94%, 35.70%] | 0.00%<br>[0.00%, 0.15%] | 0.00%<br>[0.00%, 0.15%] |
|  | Null | FPR < 0.05 | 5.10%<br>[4.27%, 6.04%] | 2.17%<br>[1.63%, 2.82%] | 4.58%<br>[3.79%, 5.47%] |
|  |  | FDR < 0.05 | 0.00%<br>[0.00%, 0.15%] | 0.00%<br>[0.00%, 0.15%] | 0.00%<br>[0.00%, 0.15%] |
|  |  | FWER < 0.05 | 0.00%<br>[0.00%, 0.15%] | 0.00%<br>[0.00%, 0.15%] | 0.00%<br>[0.00%, 0.15%] |

Like a Volcano Plot^22^, NaRnEA-based enrichment of TS-regulons for each tumor cohort can be effectively visualized by plotting the absolute value of the NaRnEA NES vs. the NaRnEA PES (Figure 2).

**Figure 2.**
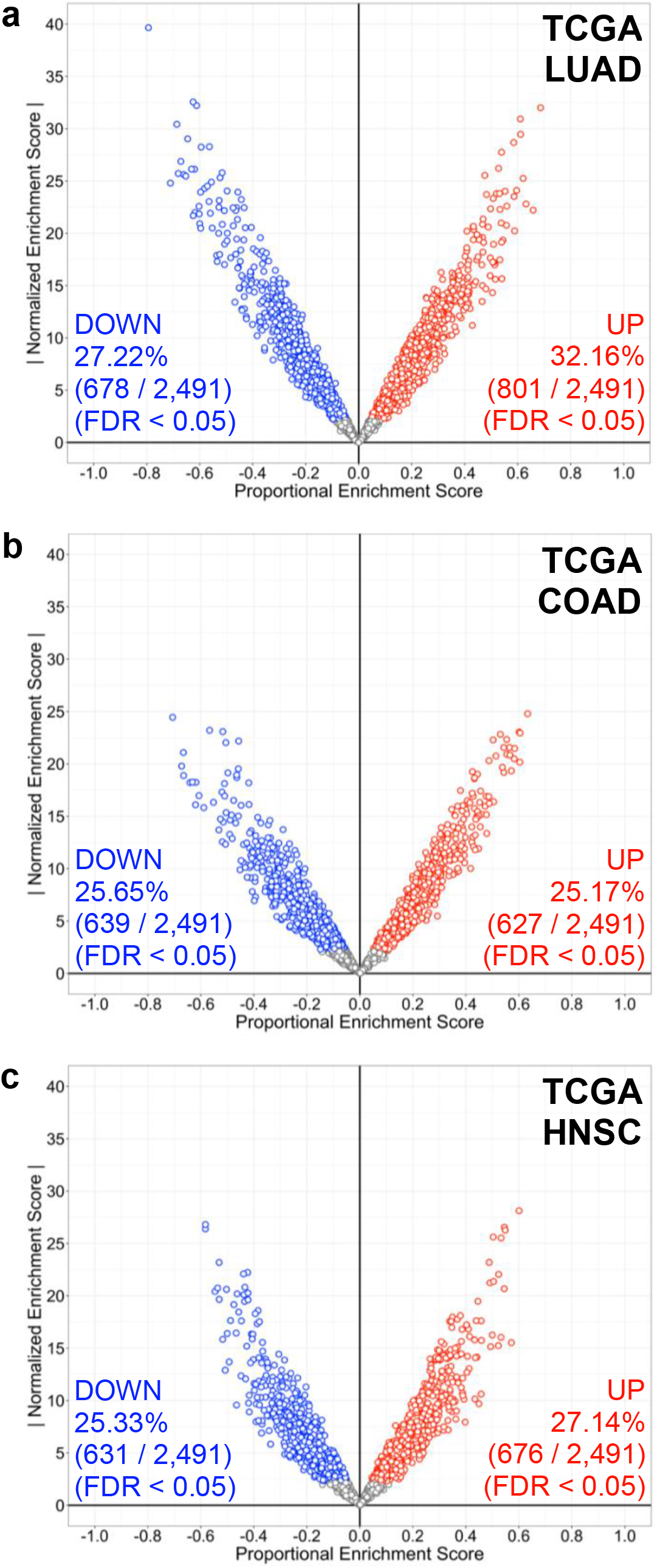
Visualizing NaRnEA-based Measurement of Differential Protein Activity in TCGA Cohorts. Visualizing NaRnEA-based differential protein activity in TCGA transcriptomic data from the (a) LUAD, (b) COAD, and (c) HNSC cancer subtypes. The absolute value of the NaRnEA Normalized Enrichment Score (NES) is plotted vs. the NaRnEA Proportional Enrichment Score (PES) to enable simultaneous inspection of the statistical significance and effect size of the enrichment, respectively. NaRnEA two-sided p-values are corrected for multiple hypothesis testing according to the methodology of Benjamini and Hochberg to control the False Discovery Rate (FDR < 0.05). The proportion of deactivated (DOWN) and activated (UP) transcriptional regulatory proteins in each of the TCGA datasets is displayed in blue and red, respectively, along with the fraction and total number of regulons which were analyzed.

### Comparison of Gene Set Enrichment and Differential Protein Abundance

We then proceeded to assess whether the differential protein activity measured by NaRnEA from TCGA transcriptomic data tracked with differential protein abundance, as experimentally assessed by mass spectrometry data from CPTAC. To this end, we used the Mann-Whitney U test to assess whether TF and coTF proteins were statistically significantly differentially abundant between primary tumor and phenotype-matched adjacent normal tissue samples, based on CPTAC proteomic data (FDR < 0.05). Specifically, we analyzed (*n*_tumor_ = 110, 97, 109) vs. (*n*_tissue_ = 101, 100, 64) from LUAD, COAD, and HNSC cohorts in CPTAC, respectively. While regulatory protein abundance is not a perfect proxy for its transcriptional activity – since protein activity is further determined by post-translational modifications, complex formation, and subcellular localization – we assumed that the two should be significantly correlated. Furthermore, since the statistical significance of differential protein activity and differential protein abundance were measured from gene expression data in TCGA and mass spectrometry data in CPTAC, respectively, the data are completely independent under the null hypothesis.

For each cancer type, we compared only the proteins for which a TS-regulon was available. Since mass spectrometry-based proteomic data do not provide a precise quantitative metric, we corrected the Mann-Whitney U test two-sided p-values from CPTAC for multiple hypothesis testing and classified each TF or coTF as upregulated (UP, Mann-Whitney U test rank-biserial correlation > 0, FDR < 0.05), downregulated (DOWN, Mann-Whitney U test rank-biserial correlation < 0, FDR < 0.05), or unchanged (NS, FDR ≥ 0.05). Similarly, we also corrected the NaRnEA two-sided p-values from TCGA for multiple hypothesis testing and classified each TF or CTF as activated (UP, NaRnEA PES > 0, FDR < 0.05), deactivated (DOWN, NaRnEA PES < 0, FDR < 0.05), or unchanged (NS, FDR ≥ 0.05).

We then tested for an association between NaRnEA-based differential protein activity from TCGA and Mann-Whitney U test-based differential protein abundance from CPTAC using a three-by-three contingency table for each cancer type (Table 2). We computed the agreement between the rows and columns using Kendall’s Tau B correlation coefficient, which adjusts for tied values within each of the three marginal categories; the statistical significance of this dependence was calculated with a Chi-Squared test. Based on this analysis, we find that the differential protein activity measured by NaRnEA in TCGA was statistically significantly positively correlated with the differential protein abundance from CPTAC for LUAD (Kendall’s Tau-B = 0.382, p = 3.14e-51), COAD (Kendall’s Tau-B = 0.281, p = 1.12e-17), and HNSC (Kendall’s Tau-B = 0.342, p = 3.63e-37) cancer types.

**Table 2.**
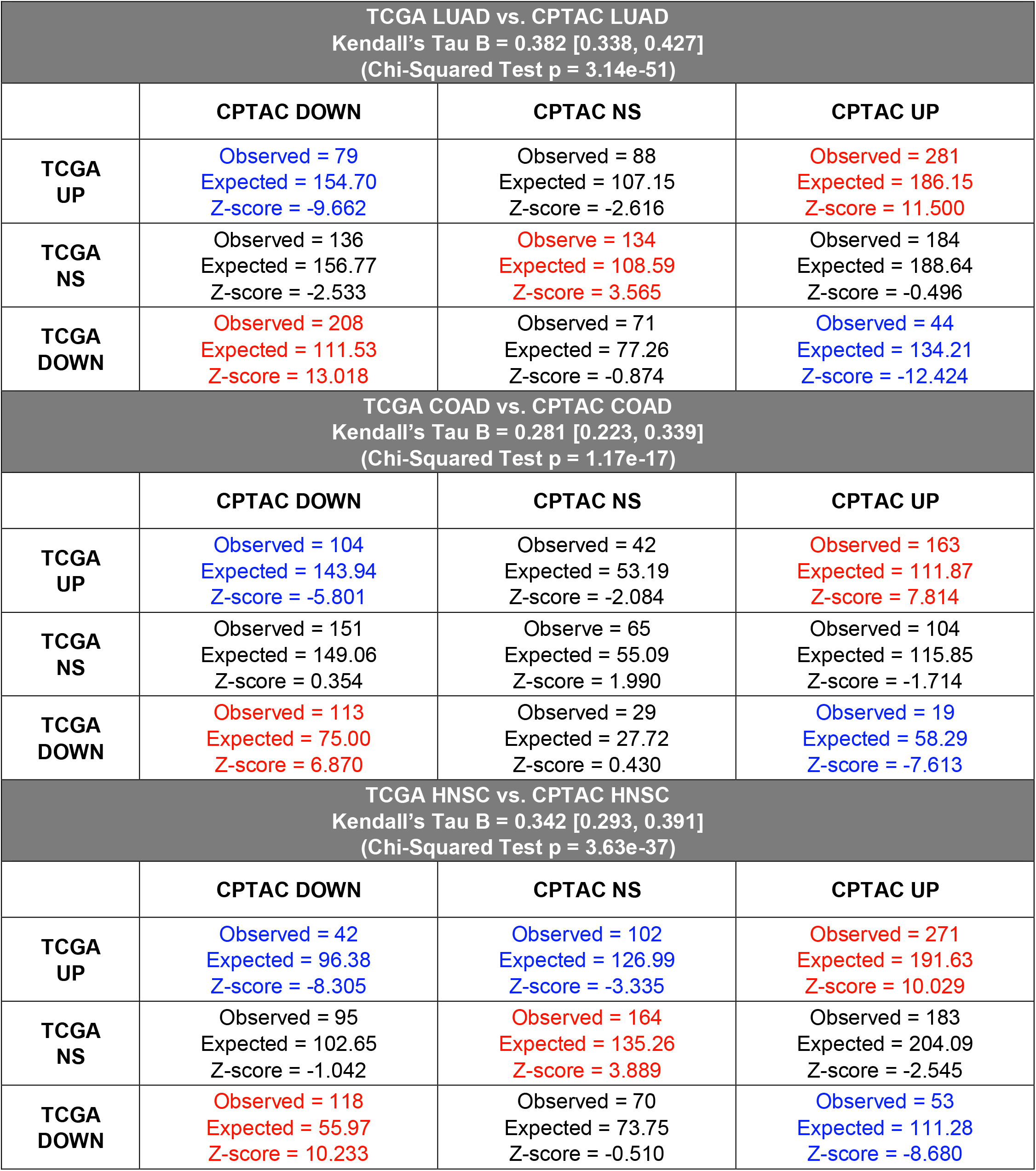
Consistency of NaRnEA-based Differential Protein Activity from TCGA with Mass Spectrometry-based Differential Protein Abundance from CPTAC. The point estimate for Kendall’s Tau-B correlation coefficient is reported along with the 95% confidence interval. Expected values for each cell are calculated under the null hypothesis of marginal independence. Z-scores are calculated by applying an inverse standard normal transformation to the quantile computed from the observed and expected values in each cell using a hypergeometric null distribution. Cells for which the z-score is significantly greater than expected under the null hypothesis (FWER < 0.05) are marked in red and cells for which the z-score is significantly less than expected under the null hypothesis (FWER < 0.05) are marked in blue.

| TCGA LUAD vs. CPTAC LUAD<br>Kendall's Tau B = 0.382 [0.338, 0.427]<br>(Chi-Squared Test p = 3.14e-51) |  |  |  |
| --- | --- | --- | --- |
|  | CPTAC DOWN | CPTAC NS | CPTAC UP |
| TCGA UP | Observed = 79<br>Expected = 154.70<br>Z-score = -9.662 | Observed = 88<br>Expected = 107.15<br>Z-score = -2.616 | Observed = 281<br>Expected = 186.15<br>Z-score = 11.500 |
| TCGA NS | Observed = 136<br>Expected = 156.77<br>Z-score = -2.533 | Observed = 134<br>Expected = 108.59<br>Z-score = 3.565 | Observed = 184<br>Expected = 188.64<br>Z-score = -0.496 |
| TCGA DOWN | Observed = 208<br>Expected = 111.53<br>Z-score = 13.018 | Observed = 71<br>Expected = 77.26<br>Z-score = -0.874 | Observed = 44<br>Expected = 134.21<br>Z-score = -12.424 |
| TCGA COAD vs. CPTAC COAD<br>Kendall's Tau B = 0.281 [0.223, 0.339]<br>(Chi-Squared Test p = 1.17e-17) |  |  |  |
|  | CPTAC DOWN | CPTAC NS | CPTAC UP |
| TCGA UP | Observed = 104<br>Expected = 143.94<br>Z-score = -5.801 | Observed = 42<br>Expected = 53.19<br>Z-score = -2.084 | Observed = 163<br>Expected = 111.87<br>Z-score = 7.814 |
| TCGA NS | Observed = 151<br>Expected = 149.06<br>Z-score = 0.354 | Observed = 65<br>Expected = 55.09<br>Z-score = 1.990 | Observed = 104<br>Expected = 115.85<br>Z-score = -1.714 |
| TCGA DOWN | Observed = 113<br>Expected = 75.00<br>Z-score = 6.870 | Observed = 29<br>Expected = 27.72<br>Z-score = 0.430 | Observed = 19<br>Expected = 58.29<br>Z-score = -7.613 |
| TCGA HNSC vs. CPTAC HNSC<br>Kendall's Tau B = 0.342 [0.293, 0.391]<br>(Chi-Squared Test p = 3.63e-37) |  |  |  |
|  | CPTAC DOWN | CPTAC NS | CPTAC UP |
| TCGA UP | Observed = 42<br>Expected = 96.38<br>Z-score = -8.305 | Observed = 102<br>Expected = 126.99<br>Z-score = -3.335 | Observed = 271<br>Expected = 191.63<br>Z-score = 10.029 |
| TCGA NS | Observed = 95<br>Expected = 102.65<br>Z-score = -1.042 | Observed = 164<br>Expected = 135.26<br>Z-score = 3.889 | Observed = 183<br>Expected = 204.09<br>Z-score = -2.545 |
| TCGA DOWN | Observed = 118<br>Expected = 55.97<br>Z-score = 10.233 | Observed = 70<br>Expected = 73.75<br>Z-score = -0.510 | Observed = 53<br>Expected = 111.28<br>Z-score = -8.680 |

After relaxing the threshold for statistically significant differential protein activity (FDR < 0.25), the same analysis demonstrated that differential protein activity measured by aREA in the TCGA LUAD cohort was statistically significantly positively correlated with the differential protein abundance from the CPTAC LUAD cohort yet with much lower statistical significance and a smaller effect size (Kendall’s Tau-B = 0.133, p = 2.75e-6). Neither the original analysis (FDR < 0.05) nor the modified analysis (FDR < 0.25) could be performed for GSEA or aREA in any of the other cohorts as no statistically significant differential protein activity could be measured by these gene set analysis methods even at the more relaxed levels. This reflects the fact that most approaches use GSEA or aREA without correcting for multiple hypothesis testing.

The low sensitivity of both GSEA and aREA can be traced directly to their reliance on null models for gene set enrichment that are estimated by an empirical phenotype-based permutation procedure this biased and underpowered. Indeed, we can show that >90% of null gene expression signatures produced by the empirical phenotype-based permutation procedure are statistically significantly correlated with the original gene expression signature. Thus, the resulting empirical null model for gene set enrichment is contaminated with enrichment scores that follow the alternative distribution rather than the null distribution, since the gene set in question will be truly enriched in the supposed null gene expression signatures which are correlated with the true gene expression signature. This results in overestimation of empirical two-sided p-values and a substantial loss of statistical power for gene set analysis.

While aREA claims to provide an alternative method for assessing the statistical significance of gene set enrichment using an analytical null model, no formal proof for the validity of this analytical null model has been published. We can show that the aREA analytical null model is also flawed by testing for the enrichment of each of the TS-regulons from the TCGA LUAD cohort in null signatures produced by shuffling gene identifiers for the LUAD tumor vs. tissue gene expression signature; these null signatures contain realistic differential gene expression values but are no longer correlated with the original gene expression signature. The resulting aREA normalized enrichment scores should be normally distributed with a mean of zero and variance of one. Using a one-sample Kolmogorov-Smirnov test we can evaluate the standard normality of the aREA normalized enrichment scores for each regulon across 1,000 shuffled gene expression signatures; after correcting for multiple hypothesis testing (FWER < 0.05), we reject the null hypothesis of standard normality for 100% of the TS-regulons. An identical analysis with NaRnEA NES values does not lead us to reject the null hypothesis of standard normality for any of the TS-regulons. This numerically confirms the mathematical accuracy of the optimal NaRnEA analytical null model while simultaneously demonstrating that the aREA analytical null model is mathematically invalid.

GSEA offers an alternative method for assessing the statistical significance of gene set enrichment using a null model estimated by an empirical gene-based permutation procedure; this approach may be regarded as an approximation of the NaRnEA analytical null model which still does not overcome the other limitations of GSEA (e.g. inability to incorporate Association Weight and/or Association Mode parameters, empirical minimum p-values determined by the number of permutations for the empirical null model, high computational complexity). Most importantly, assessing statistical significance below the 10^−5^ threshold would require an extraordinarily large number of permutations for the empirical null model, thus making the procedure computationally intractable.

Taken together, these results show that NaRnEA is dramatically more effective than both GSEA and aREA in identifying biologically relevant enrichment of gene sets.

## Discussion

It is widely appreciated that the rigor and reproducibility of scientific research depends on the use of computational and experimental methodologies which are sufficiently sensitive to make meaningful inferences while maintaining strong control over the Type I error rate to reduce spurious findings^23^. Gene set analysis methods are finding increasing application for hypothesis generation^24^, precision oncology^15^, systems pharmacology^25^, analysis of single-cell transcriptomic data^18,26^, and biomarker development^27^. Here we demonstrate that NaRnEA significantly outperforms GSEA and aREA for the purpose of gene set analysis in three independent cancer types; despite the widespread use of both competing gene set analysis methods, NaRnEA is the only method capable of consistently distinguishing between biologically coherent gene sets and gene sets constructed at random in these cohorts. Furthermore, the substantial agreement between NaRnEA-based differential protein activity in TCGA cohorts and mass spectrometry-based differential protein abundance in phenotype-matched CPTAC cohorts confirms that gene set enrichment inferred by NaRnEA cannot be explained away as erroneous false positives. Indeed, the specificity of NaRnEA is established by the fact that NaRnEA does not measure statistically significant differential protein activity for any NM-regulons in TCGA cohorts. To encourage immediate use by members of the scientific community, both NaRnEA and ARACNe3 are freely available for research use on GitHub (https://github.com/califano-lab/NaRnEA).

## Methods

### NaRnEA Mathematical Framework

The following section serves as a comprehensive derivation of the mathematical and conceptual framework underlying NaRnEA, which is designed to test for the enrichment of a gene set in a gene expression signature using a null model for gene set enrichment derived under the Principle of Maximum Entropy from information theory^8,28^; this formulation renders the NaRnEA analytical null model unique and optimal from an information theoretic perspective in the sense that any other null model must make operate under assumptions that cannot be guaranteed when the null hypothesis is true (i.e. that the gene set being analyzed is not enriched in the gene expression signature in question).

Let {*Z_g_*}*_AB_* be a vector of length (*G*), where *g* ∈ {1, …, *G*}, that represents the differential gene expression signature between biological phenotype (*A*) and biological phenotype (*B*). Let (*X_gA_*) be a random variable that represents the relative molar concentration of the transcripts originating from the *gth* gene in biological phenotype (*A*). Similarly, let (*x_gB_*) be a random variable which represents the relative molar concentration of the transcripts originating from the *gth* gene in biological phenotype (*B*). NaRnEA assumes that (*Z_gAB_*) has the following properties:

i. 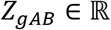
ii. *Z_gAB_* = −*Z_gBA_*
iii. 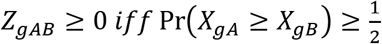
iv. 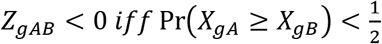
v. 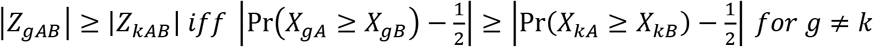

where 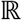 is the set of all real numbers and Pr (·) is the probability of an event.

NaRnEA tests for gene set enrichment after nonparametrically transforming the original gene expression signature; the properties of (*Z_gAB_*) are necessary and sufficient to guarantee that this nonparametric transformation preserves the relative magnitude and directionality of differential expression for each gene between phenotype (*A*) and phenotype (*B*).

NaRnEA leverages the Association Weight and Association Mode parameters so that users may construct gene sets whose members contribute unequally to gene set enrichment and which exhibit complex and/or multimodal enrichment in the gene expression signature. ARACNe3-inferred regulon gene sets for transcriptional regulatory proteins are an excellent example of such complex gene sets since the gene expression of transcriptional targets may be positively or negatively correlated with the transcriptional regulator’s activity and the expression of transcriptional targets can demonstrate varying degrees of dependency with respect to the transcriptional regulatory protein activity.

We describe the statistical framework of NaRnEA from the biological perspective of measuring differential activity of transcriptional regulatory proteins using regulon gene sets, but we note that NaRnEA can be immediately generalized to gene sets whose members share a common gene ontology or belong to the same biochemical pathway; for these gene sets, Association Weight and Association Mode parameters must be estimated based on domain-specific prior knowledge.

Let (*AW_gr_*) be the Association Weight between the *gth* gene and the *rth* transcriptional regulator. Let (*X_g_*) be the random variable representing the relative molar concentration of the transcripts originating from the *gth* gene. Let (*Y_r_*) be the random variable representing the biochemical activity of the *rth* transcriptional regulator. Let *I*[*X*; *Y*] be the mutual information between the random variables (*X*) and (*Y*). NaRnEA assumes that (*AW_gr_*) has the following properties:

i. *AW_gr_* ≥ 0
ii. *AW_gr_* > 0 *iff I*[*X_g_*; *Y_r_*] > 0
iii. *AW_gr_* ≥ *AW_kr_ iff I*[*X_g_*; *Y_r_*] ≥ *I*[*X_k_*; *Y_r_*] for *g* ≠ *k*

Let (*AM_gr_*) be the Association Mode between the *gth* gene and *rth* transcriptional regulator. Let *SCC*[*X, Y*] be the Spearman correlation coefficient between the random variables (*X*) and (*Y*). NaRnEA assumes that (*AM_gr_*) has the following properties:

i. *AM_gr_* ∈ [−1, 1]
ii. *AM_gr_* ≥ 0 *iff SCC*[*X_g_, Y_r_*] ≥ 0
iii. *AM_gr_* < 0 *iff SCC*[*X_g_*, *Y_r_*] < 0
iv. |*AM_gr_*| ≥ |*AM_kr_* | *iff* |*SCC*[*X_g_*, *Y_r_*]| ≥ |*SCC*[*X_k_, Y_r_*] | *for g* ≠ *k*

To test for the enrichment of the regulon gene set belonging to the *rth* transcriptional regulator in the gene expression signature {*Z_g_*}_*AB*_, NaRnEA calculates the Directed Enrichment Score (*D_rAB_*), which is sensitive to direction and magnitude of gene set member differential gene expression, as well as the Undirected Enrichment Score (*U_rAB_*), which is sensitive only to the magnitude of gene set member differential gene expression, as follows:

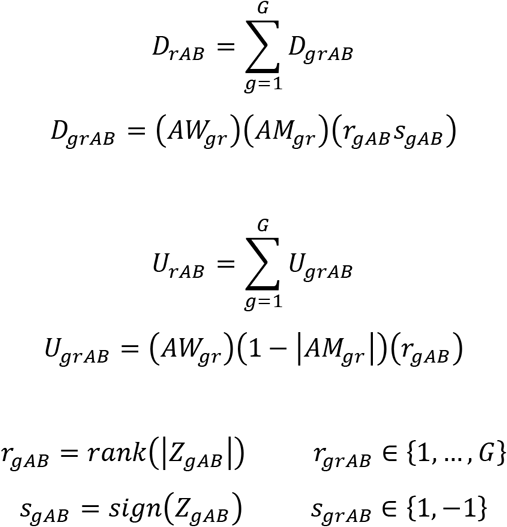

The nonparametric transformation of the original gene expression signature {*Z_g_*}_*AB*_ ↦ {*r_g_s_g_*}_*AB*_ preserves the relative magnitude and directionality of differential expression for each gene between the two phenotypes. Additionally, it simplifies subsequent calculations and allows NaRnEA to analyze gene expression signatures estimated with a variety of differential gene expression methods.

From the previously stated assumptions made by NaRnEA about the biological interpretability of (*Z_gAB_*), (*AW_gr_*), and (*AM_gr_*), we can clarify the inference of gene set enrichment using the language of frequentist null hypothesis significance testing as follows:

*H_o_*: the *rth* gene set is not enriched in {*Z_g_*}_*AB*_.
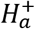: the *rth* gene set is positively enriched in {*Z_g_*}_*AB*_.
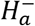: the *rth* gene set is negatively enriched in {*Z_g_*}_*AB*_.

Here the alternative hypothesis of gene set enrichment has been subdivided into two mutually exclusive versions; each of these versions implies that the members of the *rth* gene set are enriched at one or both extremes of the gene expression signature {*Z_g_*}_*AB*_. Under 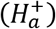 we would expect (*Z_gAB_* ≥ 0) for those members of the *rth* gene set for which (*AM_gr_* ≥ 0) and (*Z_gAB_* < 0) for those members of the *rth* gene set for which (*AM_gr_* < 0). In contrast, under 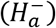 we would expect (*Z_gAB_* < 0) for those members of the *rth* gene set for which (*AM_gr_* ≥ 0) and (*Z_gAB_* ≥ 0) for those members of the *rth* gene set for which (*AM_gr_* < 0).

We can derive the marginal sampling distributions for (*D_rAB_*) and (*U_rAB_*) under (*H_o_*) using the information theoretic Principle of Maximum Entropy, which states that we should select, as our null model, the probability distribution with the greatest entropy constrained on all information available under (*H_o_*). In terms of available information under (*H_o_*), we know the following:

i. the genes for which (*AW_gr_* > 0) are members of the *rth* gene set.
ii. the members of the *rth* gene set are present in {*Z_g_*}_*AB*_.

Following the nonparametric transformation of the gene expression signature, we know that each member of the gene set may occupy one of (*G*) possible values (*r*_1_*s*_1_, …, *r_G_s_G_*}*_AB_*. Thus, for all members of the *rth* gene set, (*D_grAB_*) and (*U_grAB_*) are discrete random variables under (*H_o_*).

Let *H*[*X*] be the Shannon entropy for the random variable (*X*). Without any additional information and for (*G*) much larger than the number of members in the *rth* gene set, *H*[*D_grAB_*|*H_o_*] and *H*[*U_grAB_*|*H_o_*] are maximized when *p*(*r_gAB_s_gAB_*|*H_o_*) takes the form of a discrete uniform distribution over the entire set of (*r*_1_*s*_1_, …, *r_G_s_G_*}*_AB_* (i.e. it is equiprobable that a gene set member is located in any position in the nonparametric gene expression signature). Furthermore, without any constraints regarding higher-order dependencies between the members of the *rth* gene set under (*H_o_*), *H*[*D_rAB_*|*H_o_*] and *H*[*U_rAB_*|*H_o_*] are maximized when all members of the *rth* gene set are independent under (*H_o_*).

From this maximum entropy null model for gene set enrichment, we may derive the expected value and variance of (*D_rAB_*) under (*H_o_*) as follows:

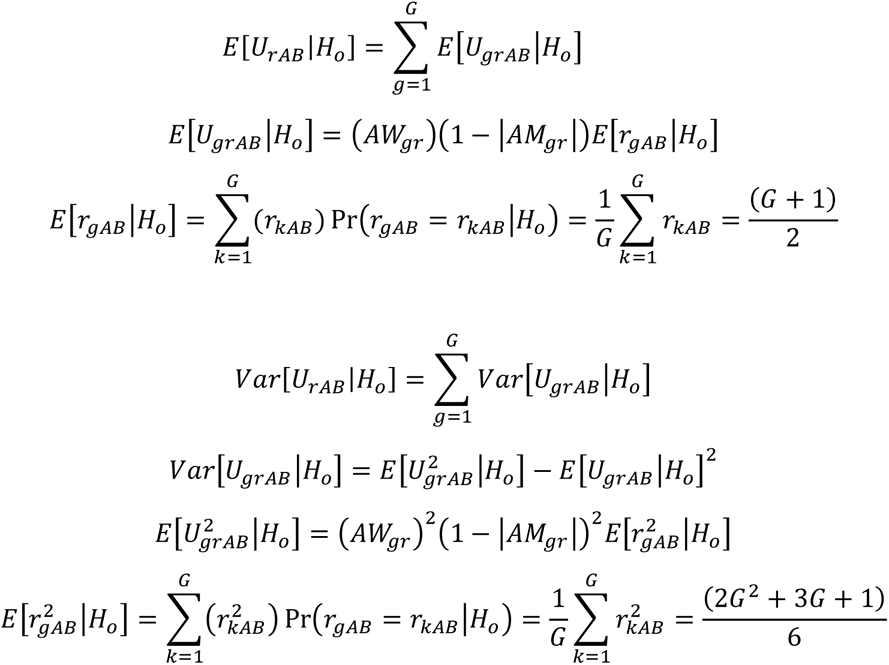

From these first two moments we can compute *NES*[*D_rAB_*], the Normalized Directed Enrichment Score for the *rth* gene set in the gene expression signature {*Z_g_*}*_AB_*, as follows:

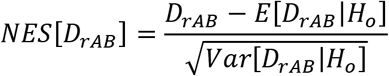

Similarly, from the maximum entropy null model for gene set enrichment, we may derive the expected value and variance of (*U_rAB_*) under (*H_o_*) as follows:

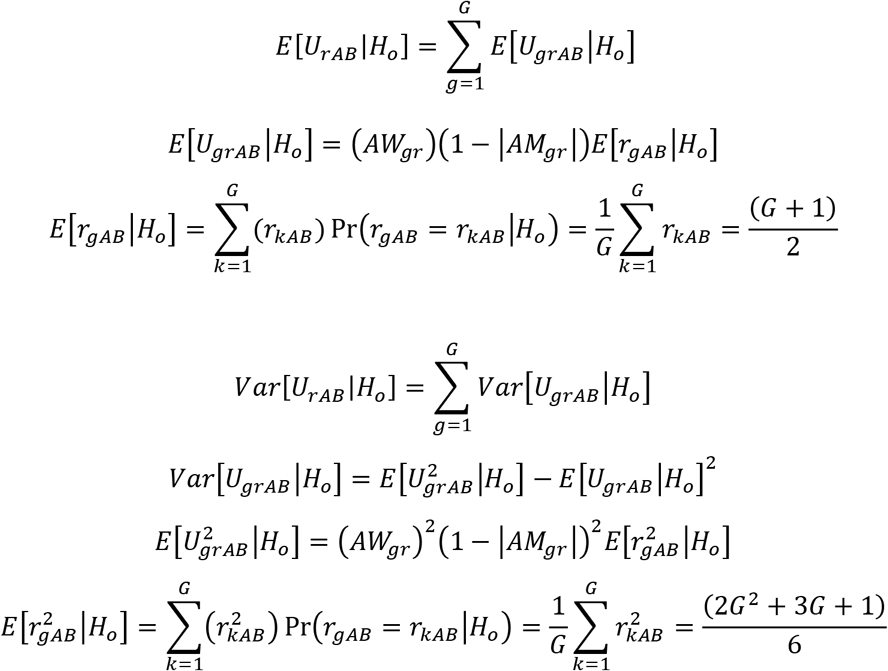

From these first two moments we can compute *NES*[*U_rAB_*], the Normalized Undirected Enrichment Score for the *rth* gene set in the gene expression signature {*Z_g_*}*_AB_*, as follows:

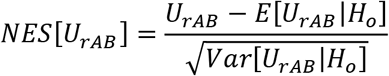

Since *NES*[*D_rAB_*] is a random variable that is equal to the sum of independent random variables under (*H_o_*), according to the Lindeberg Central Limit Theorem^29^, if

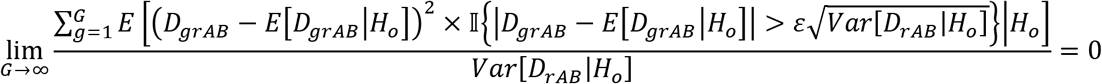

for all (*ε* > 0) where 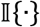 is the identity operator, then

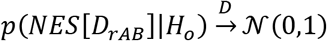

where 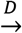 is convergence in distribution. Thus *NES*[*D_rAB_*] is asymptotically a standard normal random variable under (*H_o_*) according to the maximum entropy null model for gene set enrichment if Lindeberg’s condition is met.

Similarly, since *NES*[*U_rAB_*] is a random variable that is equal to the sum of independent random variables under (*H_o_*), according to the Lindeberg Central Limit Theorem, if

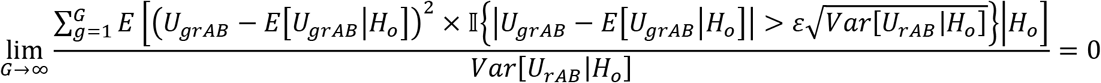

for all (*ε* > 0) where 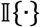 is the identity operator, then

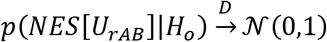

where 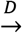 is convergence in distribution. Thus *NES*[*U_rAB_*] is asymptotically a standard normal random variable under (*H_o_*) according to the maximum entropy null model for gene set enrichment if Lindeberg’s condition is met.

Due to the nonparametric transformation of the gene expression signature, we find that this convergence to normality occurs sufficiently well to achieve strong control of the Type I error rate for gene sets with at least 30 members that are well-balanced (i.e. the Association Weight and Association Mode values for the gene set members do not vary over a large range).

Furthermore, we may generalize these statements using the multivariate central limit theorem and make the following claim about the joint distribution of *NES*[*D_rAB_*] and *NES*[*U_rAB_*] under (*H_o_*):

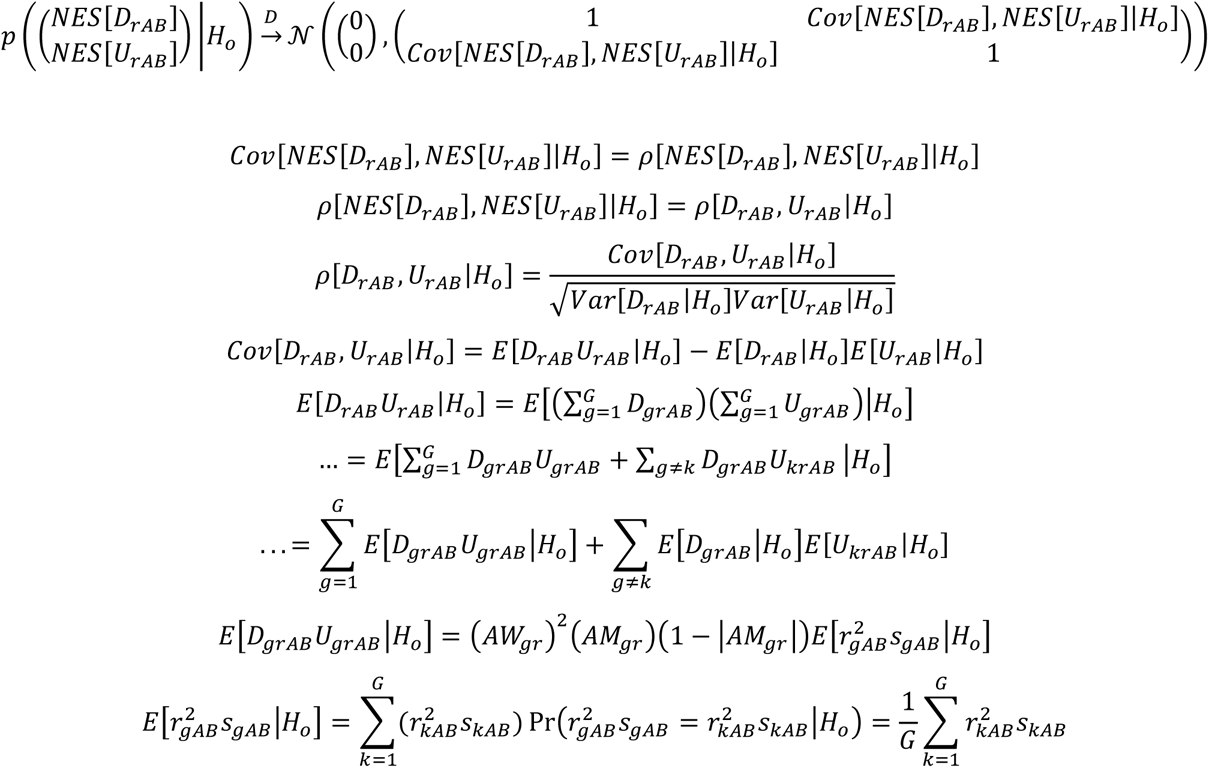

where *Cov*[*X, Y*] is the covariance between the random variables (*X*) and (*Y*) and *ρ*[*X,Y*] is the Pearson product-moment correlation between the random variables (*X*) and (*Y*).

To summarize, according to the maximum entropy null model for gene set enrichment and the multivariate generalization of the Lindeberg Central Limit Theorem, *NES*[*D_rAB_*] and *NES*[*U_rAB_*] form an asymptotically bivariate normal random vector under (*H_o_*) if Lindeberg’s condition is met for both *NES*[*D_rAB_*] and *NES*[*U_rAB_*] under (*H_o_*).

From the assumptions about the distribution of the gene set members in the gene expression signature under the alternative hypothesis, we expect (*NES*[*D_rAB_*] > 0) under 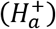 and (*NES*[*D_rAB_*] < 0) under 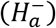. In contrast, we expect (*NES*[*U_rAB_*] > 0) under 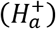 and (*NES*[*U_rAB_*] > 0) under 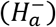. To integrate the statistical significance of *NES*[*D_rAB_*] and *NES*[*U_rAB_*] we perform the following change of variables:

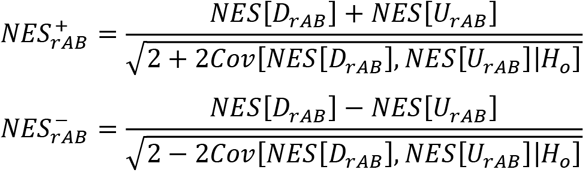

This change of variables can be represented with an affine transformation of the original bivariate normal random vector as follows:

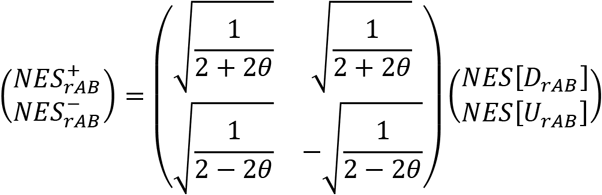

where *θ* = *Cov*[*NES*[*D_rAB_*], *NES*[*U_rAB_*]|*H_o_*].

It is a well-established result that an affine transformation of a multivariate normal random vector produces a multivariate normal random vector^30^. More formally, if (*Y* = *c* + *BX*) is an affine transformation of 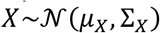 where (*c*) is a (2× 1) vector of constants and (*B*) is a (2× 2) matrix, then 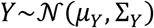, where

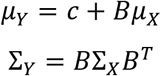

It follows from this statement that 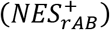 and 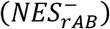 form a bivariate normal random vector under (*H_o_*) whose mean vector and variance-covariance matrix may be calculated as follows:

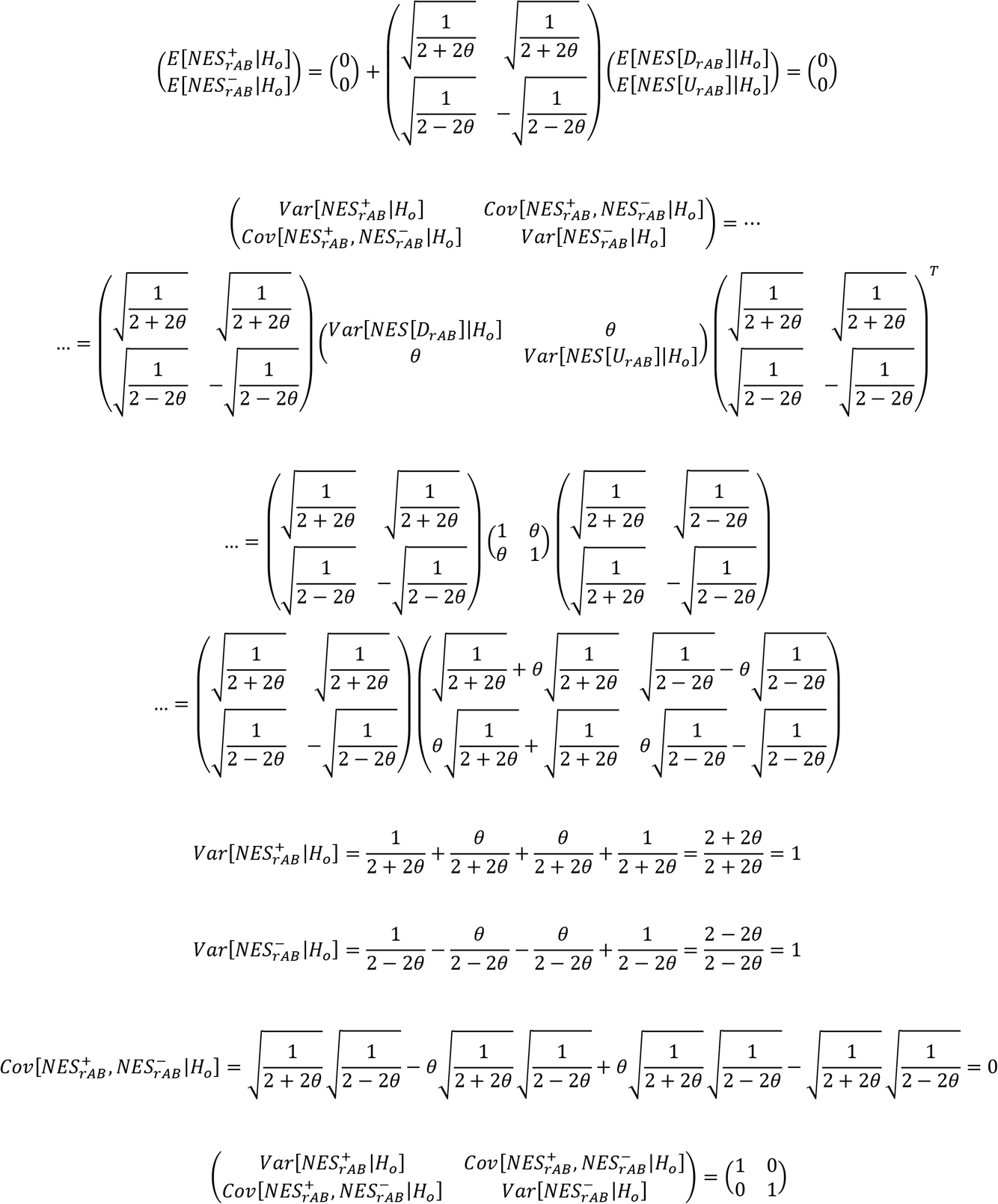

Thus, 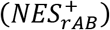 and 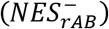 are independent standard normal random variables under (*H_o_*). This is statistically advantageous since we expect 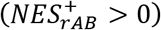 under (*H_o_*) and 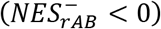 under (*H_o_*). We may leverage the asymptotic standard normality of 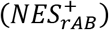 and 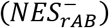 to calculate the one-tailed p-values for these test statistics as follows:

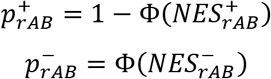

where Φ(·) is the standard normal cumulative distribution function.

Since 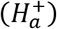 and 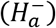 are mutually exclusive, we may use 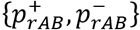 to determine which version of the alternative hypothesis is more probable. This decision constitutes a form of multiple hypothesis testing; as a result, we must adjust the final two-sided p-value to control the NaRnEA Type I error rate. Given that 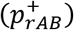 and 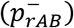 are independent under (*H_o_*), the final NaRnEA two-sided p-value for the *rth* gene set may be calculated as follows:

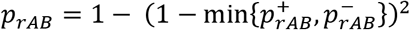

The final NaRnEA two-sided p-value is equal to the probability of observing at least one value less than or equal to 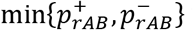 when drawing two independently and identically distributed observations from *U*[0,l], (i.e. the sampling distribution of the independent one-tailed p-values under the null hypothesis). We combine the directional information implied by 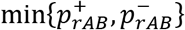 with the magnitude of the final NaRnEA two-sided p-value to calculate (*NES_rAB_*), the final NaRnEA Normalized Enrichment Score for the enrichment of the *rth* gene set in the gene expression signature {*Z_g_*}*_AB_*, as follows:

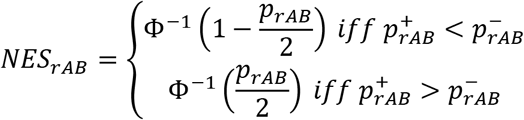

As a result, the final NaRnEA Normalized Enrichment Score for the enrichment of the *rth* gene set in the gene expression signature {*Z_g_*}*_AB_* is endowed with the following properties:

i. 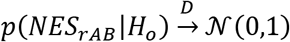
ii. 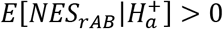
iii. 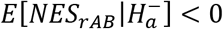

Thus, (*NES_rAB_*) serves as a single test statistic for gene set enrichment and the statistical significance of (*NES_rAB_*) may be computed analytically using the standard normal cumulative distribution function.

As a test statistic, we expect the sampling distribution of (*NES_rAB_*) under (*H_a_*) to vary depending on gene set size, parameterization, and features specific to the gene expression signature under investigation; therefore, we also require an effect size for gene set enrichment.

The NaRnEA Proportional Enrichment Score (*PES_rAB_*) for the enrichment of the *rth* gene set in the gene expression signature {*Z_g_*}*_AB_* is calculated from (*NES_rAB_*) as follows:

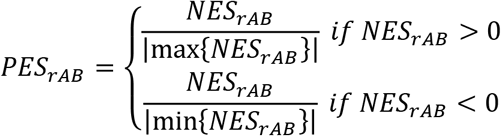

This formulation endows (*PES_rAB_*) with the following properties:

i. (*PES_rAB_*) ∈ [−1, 1]
ii. 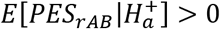
iii. 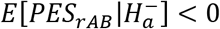

The NaRnEA Proportional Enrichment Score (*PES_rAB_*) for the enrichment of the *rth* gene set in the gene expression signature {*Z_g_*}*_AB_* functions like a correlation coefficient in that it is bounded on the interval [−1, 1], has a positive expected value when the gene set is positively enriched, has a negative expected value when the gene set is negatively enriched, and may be used to quantitatively compare the enrichment of gene sets with different parameterizations in the same gene expression signature or across different gene expression signatures. Confidence intervals for the NaRnEA Proportional Enrichment Score (*PES_rAB_*) may be approximated by bootstrapping the members of the *rth* gene set, recomputing (*PES_rAB_*) for the bootstrapped versions of the gene set, and applying an appropriate estimation procedure^31^.

Finally, NaRnEA offers post-hoc leading-edge (ledge) analysis to quantitatively evaluate the extent to which each gene set member contributes to the enrichment of the gene set in the gene expression signature. This ledge analysis is conceptually similar to the leading-edge analysis proposed for GSEA by (*Subramanian et al. 2005 Proc Natl Acad Sci*)^9^, but rather than returning a dichotomous leading edge the NaRnEA ledge analysis returns a post-hoc ledge p-value for each member of the gene set.

The NaRnEA Ledge Score for the *gth* gene with respect to the *rth* gene set in the gene expression signature {*Z_g_*}*_AB_* is computed as follows:

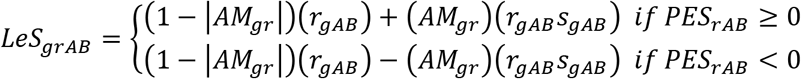

The statistical significance of (*LeS_grAB_*) may be calculated by extending the maximum entropy null model for gene set enrichment to the level of the *gth* gene as follows:

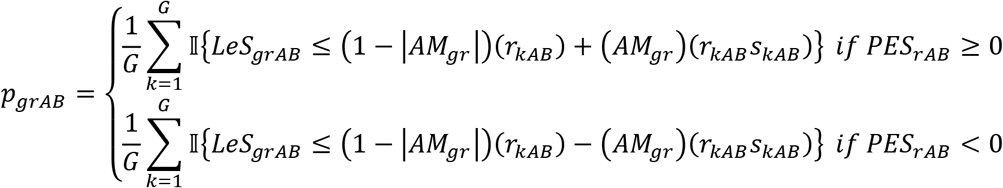

where 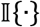 is the identity operator. The NaRnEA post-hoc ledge p-values may be corrected for multiple hypothesis testing to control the post-hoc ledge False Discovery Rate (FDR) according to the methodology of Benjamini and Hochberg^32^. The NaRnEA post-hoc ledge precision value for a gene set member is equal to one minus the minimum post-hoc ledge FDR at which the gene set member would be declared part of the gene set ledge.

### Inferring Context-Specific Transcriptional Regulatory Networks with ARACNe3

ARACNe3 is an updated implementation of the Algorithm for the Reconstruction of Accurate Cellular Networks^13,14^ which builds on the ARACNe-AP codebase^12^. The goal of ARACNe3 is to reverse-engineer a context-specific transcriptional regulatory network consisting of bivariate transcriptional regulatory interactions between a set of predefined putative transcriptional regulators and putative transcriptional targets.

ARACNe3 accepts properly normalized gene expression profiles corresponding to independent samples from a single biological phenotype. Like previous versions of the algorithm, ARACNe3 recommends that users reverse-engineer multiple estimates of the transcriptional regulatory network topology and integrate these to form an ensemble transcriptional regulatory network. However, ARACNe-AP originally recommended that the estimates of the transcriptional regulatory network topology should be reverse-engineered in a decorrelated manner by sampling from the original set of gene expression profiles with replacement (i.e. bootstrapping). While this approach is commonly employed in the field of ensemble machine learning (e.g. random forest bagging^33^), we find that sampling gene expression profiles with replacement increases the bias and variance of the adaptive partitioning mutual information (APMI) estimator; the increase in bias occurs when sampling with replacement produces areas of locally high density due to replicated data points and the increase in variance occurs because these areas of locally high density occur stochastically across different iterations of the bootstrapping procedure. To overcome this, ARACNe3 generates decorrelated individual networks by sampling ~63.21% of the gene expression profiles without replacement each time; this is equal to the probability of a unique sample appearing in a single bootstrap and thus achieves the same level of decorrelation without unduly increasing the bias or variance of the APMI estimator.

ARACNe3 estimates the null distribution for mutual information by computing the mutual information between ~1,000,000 pairs of copula-transformed gene expression marginals which are shuffled so that they are explicitly independent. ARACNe3 then fits a piecewise null model to these values where an empirical cumulative distribution function is used for the body of the null model up to the 95^th^ percentile of the data and the tail of the null model is fit analytically using robust linear regression applied to logarithmically transformed tail probabilities past the 95^th^ percentile with the mblm R package^34^. The ARACNe3 piecewise null model controls the Type I error rate for the APMI estimator more accurately than the null model implemented in ARACNe-AP and allows ARACNe3 to perform the first round of individual network pruning based on control of the False Discovery Rate (FDR), resulting in a substantial gain in power over previous versions of the algorithm which recommended performing the first round of individual network pruning based on control of the Family Wise Error Rate (FWER).

The second round of ARACNe3 individual network pruning is carried out identically to previous versions of the algorithm^13^; briefly, all three-gene cliques which remain after the first round of pruning are identified and the weakest edge of the three-gene clique is removed from the network. The final set of edges which remain after both rounds of pruning constitute an individual ARACNe3-inferred network. This procedure is carried out until one of two stopping criteria are met: (1) a prespecified maximum number of individual networks have been reverse-engineered, or (2) each putative transcriptional regulator has been assigned a prespecified minimum number of unique putative transcriptional targets. The individual networks are then integrated to form an ensemble ARACNe3-inferred transcriptional regulatory network. The mutual information and Spearman correlation for each putative transcriptional regulatory interaction in the ensemble network are estimated a final time using all gene expression profiles for greater accuracy.

The ARACNe3-inferred regulon gene set for the *rth* transcriptional regulator is constructed by extracting all putative transcriptional regulatory interactions for the *rth* transcriptional regulator from the ARACNe3-inferred context-specific transcriptional regulatory network. The Association Weight values are calculated by sorting all putative target genes based on (1) the number of individual networks in which they appeared as targets of the *rth* transcriptional regulator and (2) the final estimated mutual information between the *rth* transcriptional regulator and target gene. A copula transformation is then applied so the ARACNe3-inferred regulon gene set Association Weight values are sufficiently well-balanced to meet Lindeberg’s condition and guarantee the asymptotic standard normality of the NaRnEA Normalized Enrichment Score under the null hypothesis of gene set analysis. The Association Mode values are set equal to the Spearman correlation coefficient between the *rth* transcriptional regulator and the gene set member as estimated from all gene expression profiles.

The LUAD context-specific transcriptional regulatory network was reverse-engineered with ARACNe3 from 476 unpaired primary tumor gene expression profiles using 2,491 putative transcriptional regulators. Gene expression profiles were downloaded using the TCGAbiolinks R package from Bioconductor^35^ and normalized for sequencing depth prior to network reverse-engineering (i.e. counts per million). The first round of individual network pruning was carried out with a threshold calculated to control the FDR < 0.05. Individual networks were reverse-engineered until each putative transcriptional regulator had at least 50 unique putative transcriptional targets; this was achieved after 9 iterations. The final ensemble ARACNe3-inferred transcriptional regulatory network for TCGA LUAD has 2,491 regulators, 19,351 targets, and 947,776 regulatory interactions.

The COAD context-specific transcriptional regulatory network was reverse-engineered with ARACNe3 from 437 unpaired primary tumor gene expression profiles using 2,491 putative transcriptional regulators. Gene expression profiles were downloaded using the TCGAbiolinks R package from Bioconductor and normalized for sequencing depth prior to network reverse-engineering (i.e. counts per million). The first round of individual network pruning was carried out with a threshold calculated to control the FDR < 0.05. Individual networks were reverse-engineered until each putative transcriptional regulator had at least 50 unique putative transcriptional targets; this was achieved after 7 iterations. The final ensemble ARACNe3-inferred transcriptional regulatory network for TCGA COAD has 2,491 regulators, 19,351 targets, and 646,341 regulatory interactions.

The HNSC context-specific transcriptional regulatory network was reverse-engineered with ARACNe3 from 457 unpaired primary tumor gene expression profiles using 2,491 putative transcriptional regulators. Gene expression profiles were downloaded using the TCGAbiolinks R package from Bioconductor and normalized for sequencing depth prior to network reverse-engineering (i.e. counts per million). The first round of individual network pruning was carried out with a threshold calculated to control the FDR < 0.05. Individual networks were reverse-engineered until each putative transcriptional regulator had at least 50 unique putative transcriptional targets; this was achieved after 12 iterations. The final ensemble ARACNe3-inferred transcriptional regulatory network for TCGA HNSC has 2,491 regulators, 19,351 targets, and 810,683 regulatory interactions.

### Gene Set Enrichment Analysis

Paired gene expression profiles from TCGA were normalized using a blinded DESeq2 variance stabilizing transformation prior to analysis with GSEA which was performed as described previously by (*Subramanian et al. 2005 Proc Natl Acad Sci*)^9^ using the GSEA java command line tool from the Broad Institute’s Molecular Signatures Database (http://www.gsea-msigdb.org/gsea/downloads.jsp). The GSEA null model was estimated using 1,000 sample shuffling permutations and gene set enrichment was computed with the following parameters:

*bash GSEA_4.1.0/gsea-cli.sh GSEA -res $cur.expression.file -cls $cur.phenotype.file #Tumor_versus_Tissue -gmx $cur.gene.set.file -collapse No_Collapse -mode Max_probe -norm meandiv -nperm 1000 -permute phenotype -rnd_type no_balance -scoring_scheme weighted -rpt_label $cur.output.label -metric Signal2Noise -sort real -order descending -create_gcts false -create_svgs false -include_only_symbols false -make_sets true -median false -num 100 -plot_top_x 10 -rnd_seed $sim.seed -save_rnd_lists false -set_max 5000 -set_min 15 -zip_report false -out $cur.output.dir*

### analytical Rank-based Enrichment Analysis

Paired gene expression profiles from TCGA were normalized using a blinded DESeq2 variance stabilizing transformation prior to analysis with aREA. The VIPER R package was downloaded from Bioconductor at (https://www.bioconductor.org/packages/release/bioc/html/viper.html) and was used to run aREA as described previously by (*Alvarez et al. 2016 Nat Genet*)^4^ with the functions *viper::rowTtest, viper::ttestNull*, and *viper::msviper* using default parameters. The aREA empirical null model was estimated using 1,000 sample shuffling permutations.

### CPTAC Proteomic Data Analysis

Log-ratio normalized protein abundance data for primary tumors and adjacent normal tissues were downloaded from CPTAC for the LUAD, COAD, and HNSC cancer types from (http://linkedomics.org)^36^. Data were loaded into R and differential protein abundance between primary tumors and adjacent normal tissue was estimated for each protein with a two-tailed Mann-Whitney U test. Gene name conversion was performed using the R package biomaRt from Bioconductor^37^.

### Generation and Analysis of ROC Curves

All ROC curves were generated in R using the pROC R package from CRAN^38^. The *pROC::roc.test* function was used to conduct DeLong’s test and compare pairs of AUROC values for correlated ROC curves. The statistical significance of the AUROC was computed using a one-tailed Mann-Whitney U test. The 95% confidence intervals for the AUROC were computed using DeLong’s method. The 95% confidence intervals for ROC curves were estimated using 2,000 stratified bootstraps with the *pROC::ci.coords* function.

### Plotting and Visualization

All figures created using the *ggplot2* R package from CRAN^39^.

### Statistical Analysis

P-values were corrected for multiple hypothesis testing to control the FDR according to the methodology of Benjamini and Hochberg or to control the FWER according to the methodology of Bonferroni^32^. 95% confidence intervals for the binomial test of proportions were computed using the procedure of Clopper and Pearson^40^.

## Data Availability

All gene expression profiles from TCGA are available at (https://gdc.cancer.gov/). All protein abundance profiles from CPTAC are available at (http://linkedomics.org).

## Code Availability

NaRnEA is available as an R package on GitHub at (https://github.com/califano-lab/NaRnEA). Code necessary for running ARACNe3 can be downloaded along with the NaRnEA R package on GitHub at (https://github.com/califano-lab/NaRnEA). All code used to perform the analysis and generate the figures in this manuscript can be viewed as vignettes in the NaRnEA R package on GitHub.

## Author Contributions

Aaron T. Griffin created NaRnEA, conceived of the study, wrote the NaRnEA R code, created ARACNe3, wrote ARACNe3 wrapper scripts in R, downloaded and analyzed TCGA transcriptomic data, downloaded and analyzed CPTAC proteomic data, performed statistical analysis, created the figures, and wrote the manuscript. Lukas J. Vlahos wrote ARACNe3 C++ code, created the NaRnEA R package, and edited the manuscript. Codruta Chiuzan verified the mathematical accuracy of the NaRnEA statistical framework, verified the computational accuracy of the NaRnEA and ARACNe3 code, and edited the manuscript. Andrea Califano supervised the research, edited the manuscript, and secured funding.

## Acknowledgements

This work was supported by the National Cancer Institute (NCI) Outstanding Investigator Award R35CA197745 to Andrea Califano and the NCI Cancer Target Discovery and Development Program U01CA168426 to Andrea Califano. Aaron T. Griffin was supported, in part, by the Ruth L. Kirschstein National Research Service Award (NRSA) Institutional Research Training Grant 5T32GM007367-43, and in part by the NCI Ruth L. Kirschstein National Research Service Award (NRSA) Individual Fellowship F30CA257765. The results shown here are in part based upon data generated by the TCGA Research Network (https://www.cancer.gov/tcga) and in part based on data generated by the Clinical Proteomic Tumor Analysis Consortium (NCI/NIH). The content is solely the responsibility of the authors and does not necessarily represent the official views of the National Institutes of Health.

